# No evidence that frontal eye field tDCS affects latency or accuracy of prosaccades

**DOI:** 10.1101/351304

**Authors:** Leon C. Reteig, Tomas Knapen, Floris J.F.W. Roelofs, K. Richard Ridderinkhof, Heleen A. Slagter

## Abstract

Transcranial direct current stimulation (tDCS) may be used to directly affect neural activity from outside of the skull. However, its exact physiological mechanisms remain elusive, particularly when applied to new brain areas. The frontal eye field (FEF) has rarely been targeted with tDCS, even though it plays a crucial role in control of overt and covert spatial attention. Here we investigate whether tDCS over the FEF can affect the latency and accuracy of saccadic eye movements. 26 participants performed a prosaccade task in which they made eye movements to a sudden-onset eccentric visual target (lateral saccades). After each lateral saccade, they made an eye movement back to the center (center saccades). The task was administered before, during and after anodal or cathodal tDCS over the FEF, in a randomized, double-blind, within-subject design. One previous study (Kanai et al., 2012) found that anodal tDCS over the FEF decreased the latency of saccades contralateral to the stimulated hemisphere. We did not find the same effect: neither anodal nor cathodal tDCS influenced the latency of lateral saccades. tDCS also did not affect accuracy of lateral saccades (saccade endpoint deviation and saccade endpoint variability). For center saccades, we found some differences between the anodal and cathodal sessions, but these were not consistent across analyses (latency, endpoint variability), or were already present before tDCS onset (endpoint deviation). We tried to improve on the design of Kanai et al. (2012) in several ways, including the tDCS duration and electrode montage, which could explain the discrepant results. Our findings add to a growing number of null results, which have sparked concerns that tDCS outcomes are highly variable. Future studies should aim to establish the boundary conditions for frontal eye field tDCS to be effective, in addition to increasing sample size and adding additional controls such as a sham condition. At present, we conclude that it is unclear whether eye movements or other aspects of spatial attention can be affected through tDCS of the frontal eye fields.

## Introduction

Transcranial direct current stimulation (tDCS) harbors an exciting promise: it may influence cortical excitability and plasticity (Yavari et al., 2017), yet it is relatively noninvasive and easy to apply. These properties have attracted much attention to the technique, leading to many studies that have used tDCS to better understand the relationship between brain function and behavior (Filmer et al., 2014), to facilitate learning and to enhance cognition (Coffman et al., 2014; Cohen Kadosh, 2014) and even in clinical treatment (Lefaucheur, 2016).

Several studies have tried to enhance attention using tDCS, with mixed results (Reteig et al., 2017). In this study, we applied tDCS to the frontal eye field (FEF), a central node in the spatial attention network in the brain. Since a primary function of the FEF is the control of eye movements, we used eye tracking as our measure of tDCS efficacy.

In tDCS, a small current is passed between two electrodes, at least one of which is placed on the scalp. The current flows from the anode (positively charged electrode) to the cathode (negatively charged electrode), thereby polarizing the neurons in between. The canonical effect is that anodal tDCS enhances cortical excitability by depolarizing the resting membrane potential; cathodal tDCS on the other hand typically decreases excitability by hyperpolarizing the membrane potential (Nitsche et al., 2008).

However, the exact neurophysiological mechanisms of tDCS are much more complex and involve many more physiological processes (Jackson et al., 2016; Medeiros et al., 2012). Much of what we do know stems from in vivo and in vitro animal studies, but these findings—including the anodal vs. cathodal dichotomy—are difficult to extend to human applications (Bestmann et al., 2015; Fertonani and Miniussi, 2016).

Early human studies into tDCS focused on the motor cortex, as this allowed to assess the physiological effects of tDCS with transcranial magnetic stimulation (TMS). As expected, motor-evoked potentials elicited by a TMS pulse grew larger after anodal tDCS, and smaller after cathodal tDCS (Nitsche and Paulus, 2000, 2001). Other motor behaviors may also serve as outcome measures for tDCS. For instance, cathodal tDCS over the pre-SMA failed to suppress impulsive action tendencies (partial errors), but did prevent such impulses from expressing into full manifest errors (Spieser et al., 2015). Nevertheless, these canonical tDCS effects are not always obtained, even in the motor cortex (Strube et al., 2016).

Furthermore, tDCS parameters that work well in one brain area (e.g. the motor cortex) do not necessarily generalize to other brain regions (Bestmann and Walsh, 2017; Parkin et al., 2015). It is therefore crucial that tDCS is also applied to other, non-motor brain areas, to see to what extent its effects generalize. The dorsolateral prefrontal cortex is among one of the areas that is most frequently targeted with tDCS (Santarnecchi et al., 2015), but these studies have indeed produced more mixed results than those in the motor domain (Tremblay et al., 2014). tDCS effects on many other brain regions have not been investigated at all, or only in a handful of studies, irrespective of how well-studied they might be in other fields. The frontal eye field is a prime example of such an area.

The FEF is a key area in the dorsal visual hierarchy (Corbetta and Shulman, 2002; Schall, 2009). It is closely involved with the control of eye movements (overt attention), but is also crucial for the control of covert spatial attention (Grosbras et al., 2005; Nobre et al., 2000). Much of the evidence for a causal role of the FEF comes from stimulation studies. In fact, the FEF was first discovered when Ferrier (1873) observed that microstimulation of this area in nonhuman primates elicited contralateral saccadic eye movements.

In humans, TMS of the FEF is not strong enough to directly evoke saccades, but has been shown to affect saccade preparation. The latency of saccades generally decreases when preceded by a single TMS pulse (Juan et al., 2008; Ro et al., 2002; Thickbroom et al., 1996). Repetitive and theta-burst TMS protocols on the other hand can slow saccades for a more prolonged period of time (Nyffeler et al., 2006). TMS of the FEF has also been shown to impair covert attention (Capotosto et al., 2009).

In spite of the ubiquitous role of the FEF in visuospatial attention, tDCS of the FEF is largely uncharted. In contrast to TMS, which generally seems to have an inhibitory effect, one attractive feature of (anodal) tDCS is that it could enhance FEF activity, and thereby spatial attention (Reteig et al., 2017). Kanai et al. (2012) were the first to probe for effects of anodal or cathodal FEF-tDCS on prosaccades (saccades to a target) and antisaccades (away from a target). Their main finding was that anodal tDCS decreased the latency of contralateral prosaccades (i.e. tDCS of the left FEF slows saccades to targets in the right visual hemifield, or vice versa). For antisaccades, they observed a different pattern: cathodal tDCS increased the latency of ipsilateral antisaccades, and anodal tDCS reduced the frequency of erroneous saccades to the target. They further explored whether tDCS also affects the accuracy of saccades, but found no effects on either the mean deviation or the variability of saccade endpoints.

We identified just four more studies that have attempted FEF-tDCS. Similar to the main finding in Kanai et al. (2012), Tseng et al. (2014) showed that anodal FEF-tDCS shortened the latency of prosaccades to a (neutral) face stimulus in the presence of distractors (fearful or scrambled faces). In contrast, Chen and Machado (2017) found no effects of anodal or cathodal tDCS on either pro- or antisaccades, even though their study closely resembled the one by Kanai and colleagues. Two more studies paired FEF-tDCS with a visual search task (Ball et al., 2013; Ellison et al., 2017), which is known to depend on the FEF (Reynolds and Chelazzi, 2004), but tDCS did not affect reaction times in either study. Jointly, these studies paint a mixed picture of frontal eye field tDCS efficacy.

Nevertheless, the main result of Kanai et al. (2012)—that anodal tDCS speeds contralateral prosaccades—seems plausible for several reasons. First, the behavioral enhancement following anodal tDCS (i.e. faster saccade latencies) is in accord with the general tDCS literature. Because this enhancement was specific to the contralateral hemifield, it is unlikely to be a placebo or general arousal effect. Finally, there is a clear candidate mechanism for the effect. Seminal work has shown that monkeys make a saccade as soon as the activity in the FEF reaches a certain threshold (Hanes & Schall, 1996). Assuming that anodal tDCS increases excitability of the FEF, this threshold would be reached sooner, and saccade latency would therefore decrease.

Both the sample size and the effect size in Kanai et al. (2012) were on the smaller side: anodal tDCS shortened saccade latency by around 6 ms, and there were 16 participants in each group (anodal and cathodal). Indeed, Chen and Machado (2017) did not find this effect, even though their study was highly similar. Recently, the number of tDCS studies that have produced null results has grown steadily (see the other studies in this Research Topic), thereby casting doubt on the efficacy of the technique and the replicability of the existing tDCS literature (Horvath et al., 2014, 2015; Medina and Cason, 2017).

We therefore performed a conceptual replication of Kanai et al. (2012) (see the Discussion section for a table of all the differences between the present study and theirs). Participants performed a prosaccade task before, during and after anodal or cathodal tDCS over the right FEF, in a (randomized, double blind) within-subject design. We hypothesized to find the same effect as Kanai et al. (2012)—anodal tDCS should decrease the median latency of lateral saccades to targets in the left hemifield (i.e. contralateral to the stimulated right FEF). We also conducted exploratory analyses of the full saccade latency distribution. Next to saccade latency, we also probed for effects of tDCS on the accuracy of saccades (mean deviation and variability of saccade endpoints), although Kanai et al. (2012) did not find any. Finally, in addition to Kanai et al. (2012), we also measured the saccades participants made back to the center, after each lateral saccade.

## Material and methods

### Participants

31 participants took part in the study; data from 26 participants (14 female, mean age = 25.9, range = 21-34, *SD* = 3.42) were included in the analyses (see Participant and saccade exclusion in the Results section). The experiment and recruitment took place at the University of Amsterdam; all procedures for this study were approved by the ethics committee of the Department of Psychology, and complied to relevant laws and institutional guidelines. Participation was precluded in case screening with a tDCS safety questionnaire revealed potential issues, including (a history of) neurological, psychiatric or skin conditions. All participants gave written informed consent and were compensated with course credit or money (€10 per hour).

### Procedure

The study followed a randomized, double-blind, crossover design, in which subjects received anodal and cathodal tDCS in separate sessions (Figure 1). The two sessions were separated by a washout period of at least 48 hours to minimize the risk of carry-over effects.

Neuronavigation was always performed at the start of the first session (see Frontal eye field localization), and was usually not repeated on the second session. Otherwise, the procedure for each session was identical. First, a brief trial stimulation allowed participants to experience the sensations induced by tDCS and to decide whether they wanted to continue with the experiment (see tDCS). After setting up the eye tracker (see Eye tracking), participants practiced the prosaccade task (see Task) for one block (120 trials). Subsequently, participants performed 12 blocks of the task in three phases (Figure 1): 3 blocks prior to stimulation (*baseline*), 3 blocks during stimulation (anodal or cathodal tDCS), and 6 blocks after stimulation (*post-1* and *post-2*). Each block lasted approximately 5 minutes.

During the stimulation phase, the first block started after ramp-up of the current (1 minute). If the participant finished the required 3 blocks of the task within the next 16 minutes (15 minutes of constant stimulation and 1 minute of ramp-down), they were asked to sit quietly, until the stimulation had completed.

After task performance was complete, participants filled in a questionnaire on the occurrence of adverse effects related to tDCS (see Supplementary Figure 1).

**Figure 1:**
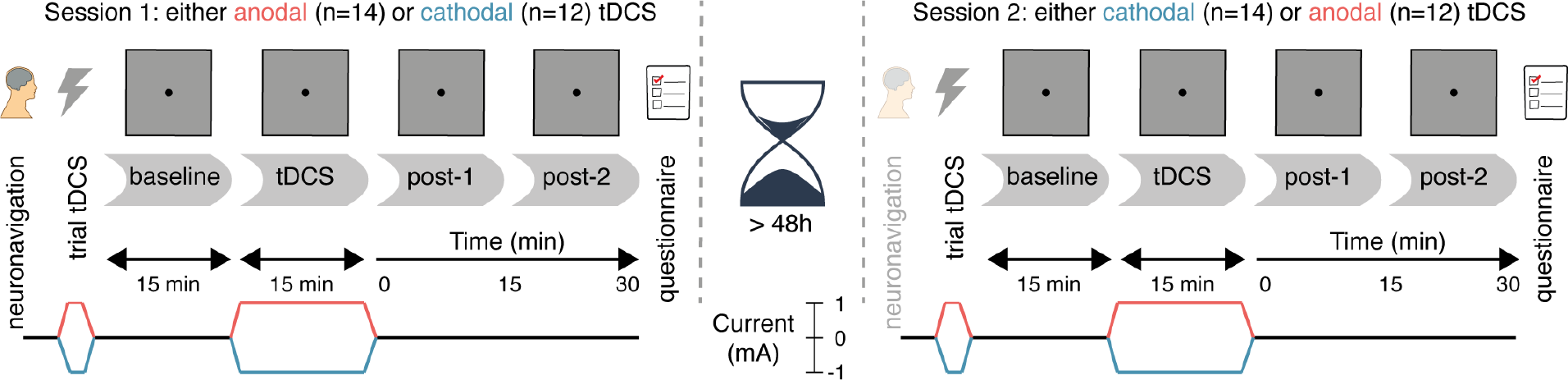
Experimental design. After a baseline measurement, participants received either anodal or cathodal tDCS while continuing to perform the prosaccade task, followed by two more post-tDCS assessments. After a washout period of at least 48 hours, the second session followed the same protocol, except that the tDCS polarity was opposite (e.g. if participants received anodal tDCS in the first session, cathodal tDCS was applied in the second session, and vice versa).

**Figure 2:**
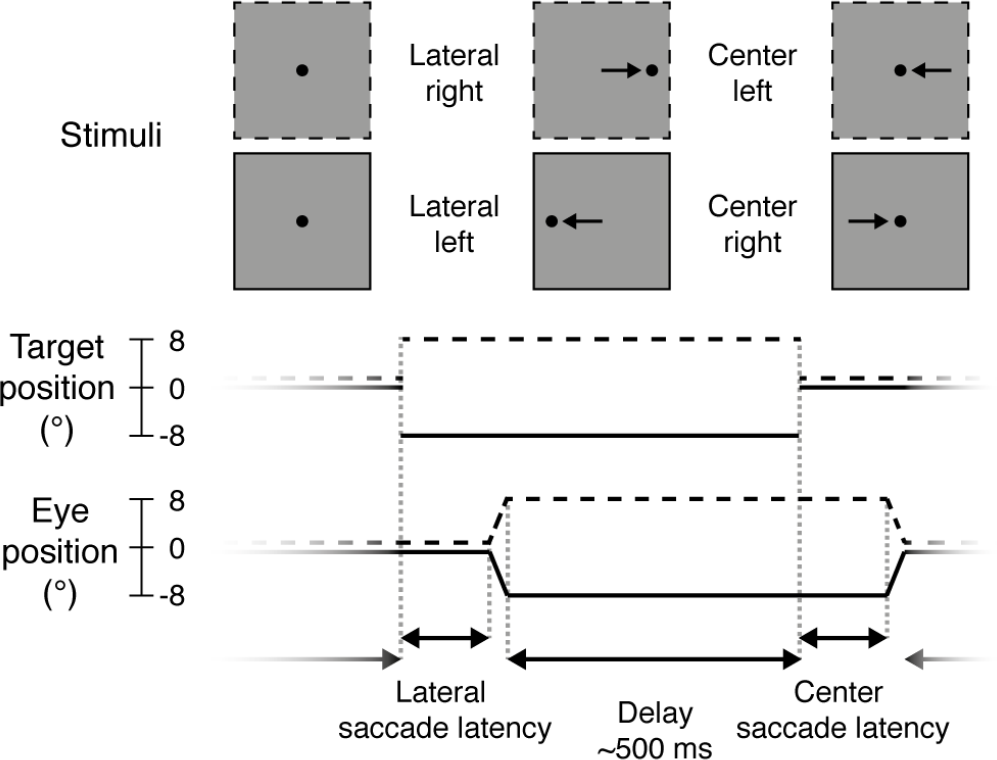
Prosaccade task. Each trial started with a *lateral saccade*, where the participant made an eye movement in response to the target stimulus (black dot) jumping from the center of the screen to either the *right* (dotted lines, +8°) or *left* (solid lines, −8°). After a delay period (mean: 500 ms) following saccade offset, the target jumped to the *center* again and participants made a *left*ward or *right*ward saccade back to it. After this saccade there was again a delay period, before the next trial started and the target appeared to the left or right again.

### Task

Participants performed a no-gap, no-overlap prosaccade task (Figure 2) similar to the task in Kanai et al. (2012), in which they had to make eye movements to a target stimulus. Stimuli were displayed using MATLAB (The MathWorks Inc.) and Psychtoolbox-3 (Brainard, 1997; Kleiner et al., 2007; Pelli, 1997).

Each trial started with the participant fixating the target (black dot, diameter: 0.5 degrees of visual angle, henceforth: °) in the center of the screen. The target would then disappear and instantly reappear to either the left or right side of the screen (8° from center), prompting the participant to make an eye movement to the new location of the target (*lateral saccade*). The target would then jump back to the center, and again the participant made an eye movement to it (*center saccade*). Each trial thus required two prosaccades: one to an unpredictable location (lateral saccades, either to the left or right) and one to a predictable location (center saccades, always back to center, so in the opposite direction as the preceding lateral saccade).

After the target appeared at a new location, saccades were monitored online for 400 ms. The target remained at that location for a variable delay period, starting from the time of the saccade endpoint. If no saccade was detected (with an accuracy within 2° from the target location), the delay period started after the saccade monitoring period ended (i.e. after 400 ms). The delay duration was drawn randomly from an exponentially decaying distribution with a mean of 0.5 seconds, truncated between 0.3 and 3 seconds.

Every 20 trials, participants could take a brief self-timed break. After a block of 120 trials, participants could take a longer break and remove their head from the eye tracker chin rest. Target location (left or right side of the screen) was pseudorandomly distributed across trials within a block.

At the end of each block, a feedback screen was presented that displayed the average accuracy (in mm) and speed (in ms) of the lateral saccades within the block. The task instruction was to make saccades as quickly and accurately as possible, but with an emphasis on speed.

### Eye tracking

The right eye position was sampled at 1000 Hz with an EyeLink 1000 (SR Research Ltd.) eye tracker. During eye tracking, a chin- and forehead rest were used to keep the head in place. The tracker was calibrated with a standard 9-point calibration before the first block and after each subsequent block. If necessary, calibration was redone until no calibration point had an error larger than 1°, and the average error was below 0.5°.

The EyeLink 1000 online parser was used to classify the raw data samples into saccades, fixations and blinks. We used the default parameters for detecting saccade on-and offsets: when the eye velocity and acceleration both crossed a threshold of 30°/s and 8000°/s2, respectively. We extracted only the first saccade that was detected after the target moved, provided it was larger than 1.5°, to exclude microsaccades made when the participant was still fixating.

### tDCS

tDCS was delivered online (i.e. during performance of the prosaccade task) using a DC-STIMULATOR PLUS (NeuroCare Group GmbH). The current was ramped up to 1 mA in 1 minute, followed by 15 minutes of stimulation at 1 mA, after which the current was ramped down again in 1 minute.

One 3×3 cm electrode (9 cm^2^, current density: 0.11 mA/cm^2^) was placed over the right frontal eye field; the other electrode was 7×5 cm (35 cm^2^, current density: 0.029 mA/cm^2^) and was placed on the left forehead, centered above the eye. The rubber electrodes were fixed to the scalp with Ten20 conductive paste (Weaver & Co.). Participants received either anodal (anode over FEF, cathode on forehead) or cathodal (cathode over FEF, anode on forehead) tDCS, in separate sessions.

Both the participant and the experimenters were blind to the polarity of the stimulation (anodal or cathodal). The experimenter loaded a stimulation setting on the tDCS device (programmed by someone not involved in this study), without knowing whether it was mapped to deliver anodal or cathodal tDCS. In the second session, the electrodes were connected to the positive and negative terminal of the device oppositely to the first session, such that the opposite polarity was applied. The participant was not informed about this difference until after the end of the second session.

Before starting the task, a trial stimulation was given after which participants were explicitly offered to terminate the experiment if the tDCS was too uncomfortable. For the trial stimulation, the current ramped up to 1 mA in 45s, stayed at 1 mA for 15s, and ramped down again in 45s. No participant opted to terminate the experiment.

### Frontal eye field localization

We localized the right frontal eye field for each participant using pre-existing MRI scans. All participants had a T1 structural scan available; for 5 participants we also used functional MRI data from a retinotopic mapping experiment (Es et al., 2017), and targeted retinotopic region sPCS (Mackey et al., 2017).

The presumed location of the FEF was defined as slightly lateral to the superior frontal sulcus, in the anterior bank of the pre-central sulcus (Amiez and Petrides, 2009; Blanke et al., 2000; Mackey et al., 2017; Vernet et al., 2014). For the retinotopic mapping data, we used the coordinate of the peak voxel in the cluster positioned closest to this location.

To obtain the MNI coordinates of the presumed FEF for each participant, we used FSL (Jenkinson et al., 2012; Smith et al., 2004) BET (Smith, 2002) to extract the brain and FLIRT (Jenkinson and Smith, 2001; Jenkinson et al., 2002)) to register it to the MNI152 template.

At the beginning of the first session, neuronavigation was performed using the visor2 system (ANT Neuro). We placed a marker in the imaged brain 5 mm posterior to the presumed FEF location, to increase the likelihood that the current would flow through the FEF from/to the forehead electrode. The location on the scalp directly above this marker (i.e. parallel to the inferior-superior axis) was stained with surgical skin ink. The tDCS electrode was then centered on this ink mark. If the ink mark was no longer visible in the second session, neuronavigation was repeated.

### Analyses

Data were analyzed using the R programming language (R Core Team, 2017) and several general packages (Robinson, 2017; Wickham, 2017; Wickham and Grolemund, 2017; Wilke, 2016) from within RStudio (RStudio Team, 2016).

#### Saccade measures

To determine the effects of frontal eye field tDCS on eye movement behavior, we examined three different measures, following Kanai et al. (2012): saccade latency, saccade endpoint deviation, and saccade endpoint variability. Saccade latency was defined as the time between the onset of the target stimulus and the onset of the saccade. We computed the median saccade latency instead of the mean, as the distribution of saccade latencies tends to be heavily right-skewed. Saccade endpoint deviation was defined as the Euclidian distance (shortest straight line) between the saccade endpoint and the actual target position. Saccade endpoint variability was defined as the standard deviation of the horizontal coordinates of the saccade endpoints.

#### Quantile analysis

To improve sensitivity, we also probed for potential differences between anodal and cathodal tDCS across the entire distribution of saccade latencies. For instance, it is conceivable that tDCS has no effect on median saccade latency, but only on very fast (or slow) saccades, as these may involve different cognitive or neurophysiological processes.

We therefore created *shift functions* (Rousselet and Wilcox, 2016; Rousselet et al., 2017) based on the saccade latency distributions for each subject and condition. In this method, the deciles of each distribution (i.e. the 9 values that split the distribution in 10 equal parts) are computed using a Harrel-Davis quantile estimator (Harrell and Davis, 1982). For each subject and condition, the deciles for the anodal and cathodal distributions were then subtracted, and 95% confidence intervals of the decile differences were computed using a percentile bootstrap (Wilcox and Erceg-Hurn, 2012). For each individual subject, significance is then assessed and corrected for the 9 decile comparisons using Hochberg’s method (Hochberg, 1988). We report the average shift function across participants and the number of subjects that show a significant difference for each decile.

#### Trial selection

Following Kanai et al. (2012), we rejected saccades when (1) eye position at saccade onset deviated from fixation (i.e. the previous target location) by more than 1.8°, (2) the saccade endpoint deviated from the target position by more than 8° (e.g. if participants made a saccade in the wrong direction), (3) saccade latency was below 50 ms, or (4) saccade latency exceeded 400 ms. We did not reject any saccades for the quantile analyses, because the tails of the saccade latency distribution were of primary interest, and fixation and saccade errors were rare (see Participant and saccade exclusion in the Results section).

The remaining trials were collapsed across three blocks within one phase of the experiment (e.g. all the blocks during tDCS) to maximize the amount of trials that went into each average. Data were therefore analyzed over four time periods: *baseline*, *tDCS*, *post-1* and *post-2*.

#### Statistics

For each saccade measure, paired-sample t-tests were run on the baseline data of each session (i.e. anodal baseline vs. cathodal baseline), to check for any differences prior to stimulation onset. Subsequently, we subtracted the average scores during the *baseline* period from the three other periods (*tDCS*, *post-1* and *post-2*), to assess the change from baseline for each individual.

Repeated measures ANOVAs were conducted (Lawrence, 2016) with the same factors as Kanai et al. (2012): Stimulation (anodal vs. cathodal), Time Period (during tDCS, post-tDCS [0-15 min], post-tDCS [15-30 min]), and Saccade Direction (left vs. right). Statistics for all main effects and interactions involving the Stimulation factor are reported in tables. We ran separate ANOVAs for lateral saccades and center saccades, because Kanai et al. (2012) did not measure the latter. Effect sizes were computed as generalized eta squared 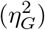 (Bakeman, 2005). Violations of the assumption of sphericity where detected with Mauchly’s test, in which case Greenhouse-Geisser corrected p-values are reported. Paired sample t-tests were conducted to follow-up on significant effects in the repeated measures ANOVA.

We also conducted Bayesian analogues of these repeated measures ANOVAs (Rouder et al., 2012, 2016) using the BayesFactor R package (Morey and Rouder, 2015) with the default prior specification. Bayes factors are reported both in terms of evidence for the alternative hypothesis (BF_10_) as well as the null hypothesis (BF_01_). We used the scheme from Wagenmakers et al. (2017) to classify the strength of the evidence (e.g. a BF from 1-3 can be considered “anecdotal” evidence, BFs 3-10 “moderate” evidence, etc.). We computed Bayes Factors comparing the null model (intercept and random effect of participant) against all other models, only excluding models containing exact cross-over interactions (i.e. interactions without the constituent main effect), to decrease the model space (Rouder et al., 2016).

Still, with 3 factors in the design, this analysis produces 19 Bayes factors, complicating model comparison (Wagenmakers et al., 2017), and comparison of the Bayesian and the classical ANOVAs. We therefore also quantified the evidence for experimental effects instead of just individual models, by computing an “inclusion Bayes factor across matched models” (concept and terminology borrowed from the JASP software package (JASP Team, 2018)). Briefly, for each effect, this Bayes Factor compares two subsets of models: (1) the subset of models that contain the effect of interest, but no higher order interactions; (2) the subset of models that result from stripping the effect of interest from (1). The inclusion Bayes factor thus reflects the evidence for an effect of interest, based not on just a single model, but on the posterior probabilities of all models that include this effect. Bayesian paired-sample t-tests were conducted to follow-up on effects with an inclusion BF higher than 10 (“strong”, “very strong”, or “extreme” evidence (Wagenmakers et al., 2017)).

### Data, materials, and code availability

All code used for this study is available on GitHub (https://github.com/lcreteig/sacc-tDCS), including R notebooks (Xie, 2015, 2016) that demonstrate how to reproduce all the results, figures and statistics from the data. The eye tracking, questionnaire and meta-data can be downloaded from a figshare repository (Reteig et al., 2018). All of these and additional resources can be found on this study’s page on the Open Science Framework (https://osf.io/8jpv9).

## Results

### Participant and saccade exclusion

Data from 26 participants were included in the analyses. 14 participants received anodal before cathodal stimulation; 12 participants received cathodal before anodal stimulation. Two participants were excluded because their two sessions were separated by less than 48 hours due to a scheduling error. Three more participants were excluded because they had fewer than 50 saccades left per cell after rejecting outlier saccades. For the remaining 26 participants, 2.0% of all saccades were rejected because they were too fast (latency < 50 ms), and 2.6% were rejected because fixation was inaccurate (deviation > 1.8°). Too slow saccades (.12%) and saccade direction errors were almost non-existent (.16%). This left an average of 175 lateral saccades (range: 142-180) and 156 center saccades (range: 74-180) per cell.

**Figure 3:**
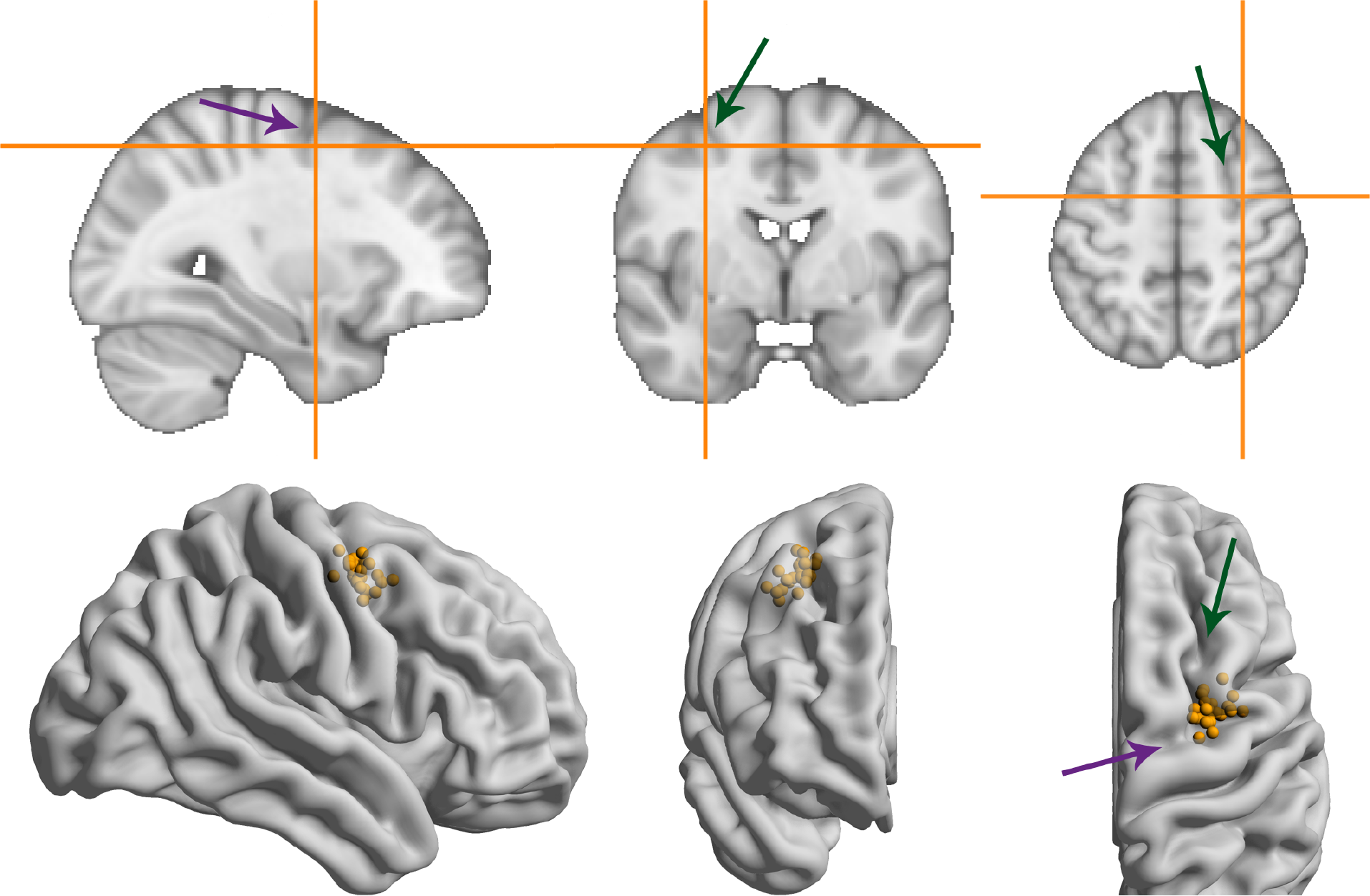
MNI coordinates of the right frontal eye field. Green/more vertical arrows indicate the superior frontal sulcus, purple/more horizontal arrows indicate the pre-central sulcus. (**A**) Average MNI coordinate across participants. (**B**) Coordinates for individual participants overlaid on a glass brain representation of the MNI template using Surf Ice software (Rorden, 2017).

### Neuronavigation

Figure 3 shows the MNI coordinates of the presumed right frontal eye field. While there is some spread (see Supplementary Table 1 for the coordinates of all participants), the average coordinate (31.5, −1.8, 51.6) matched the anatomical definition we used for the individual MRIs: slightly anterior to the pre-central sulcus and slightly lateral to the superior frontal sulcus. The average coordinate also lies close to the one used in Kanai et al. (2012), which was taken from Paus (1996) (31.3, −4.5, 50.9).

### Median saccade latency

We hypothesized that anodal tDCS would increase excitability of the frontal eye field, such that the threshold for making a saccade would be reached sooner. Specifically, we predicted a decrease in median latency of leftward saccades (contralateral to the stimulated right FEF), based on earlier findings that anodal tDCS speeded contralateral saccades by 6.4 ms compared to baseline (Kanai et al., 2012).

The latency changes in our data were more modest and did not exceed 4 ms for any condition (6). In contrast to Kanai et al. (2012), we found no effect of anodal tDCS on contralateral saccade latency, as reflected in a non-significant interaction between Stimulation and Saccade Direction for lateral saccades, and moderate evidence for the null hypothesis (Table 1). The average change from baseline for leftward lateral saccades in the anodal session were all less than 1 ms (tDCS: *M* = −0.17, *SD* = 5.34; post-1: *M* = −0.62, *SD* = 7.13; post-2: *M* = 0.96, *SD* = 9.65).

tDCS also did not seem to affect lateral saccade latency in other ways: all the effects with the factor Stimulation were non-significant and the null-hypothesis was always supported more than the alternative. From the full ANOVA for lateral saccades, the only significant effects were a main effect of Time Period (F(2,50) = 3.46, *p* = .039, 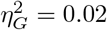) and a Time Period by Saccade Direction interaction (F(2,50) = 3.66, *p* = .033, 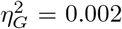).

**Figure 4:**
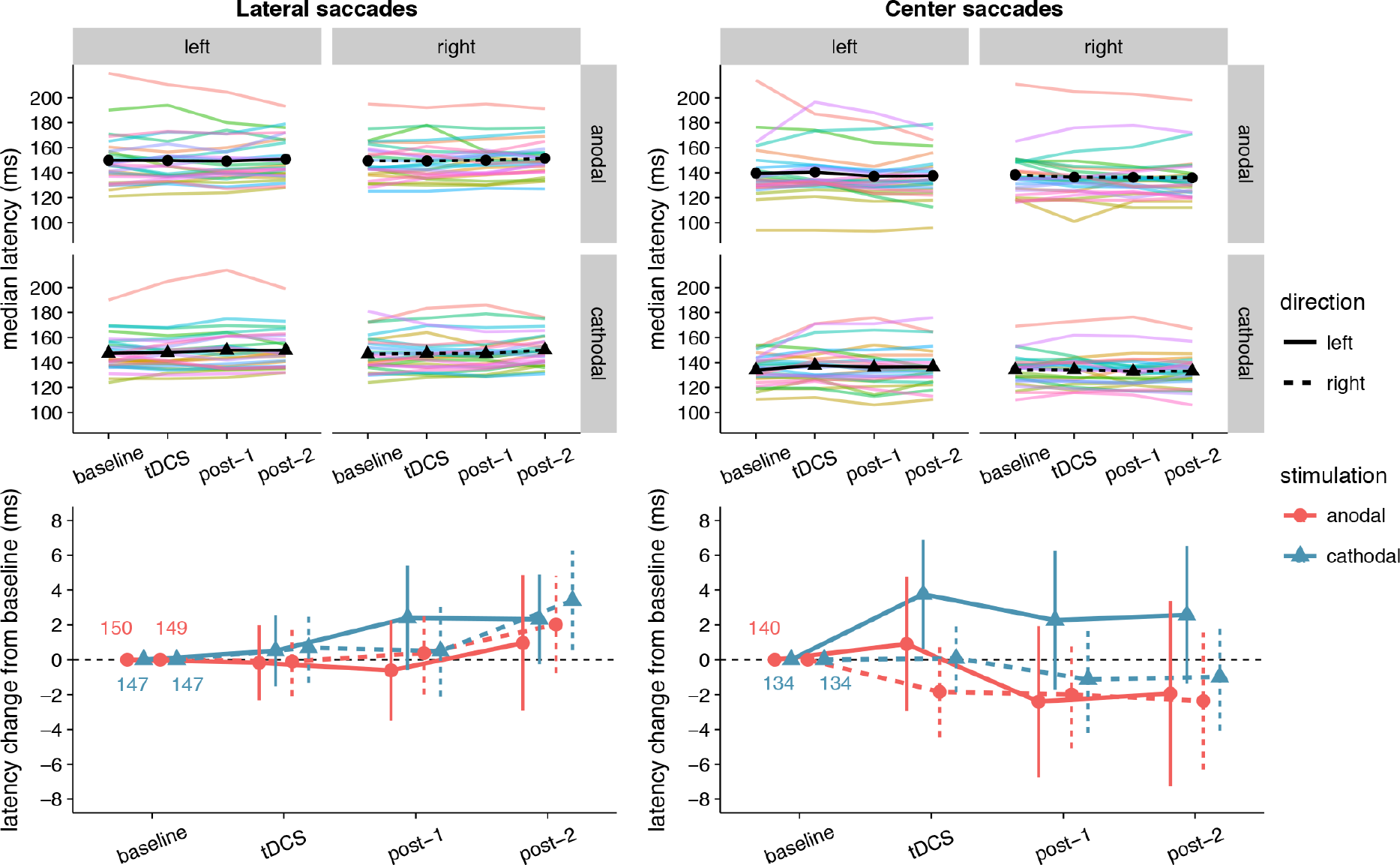
Effects of frontal eye field tDCS on saccade latency. Data are shown for left vs. rightward saccades, in the anodal vs. cathodal session, for four 15-minute time periods: baseline, during tDCS, and after tDCS (post-1 and post-2). **Top row**: Colored lines show data from individual participants; black lines show the group median. **Bottom row**: Change in saccade latency after baseline subtraction. Numbers inside the plot axes are the baseline saccade latencies per condition. Error bars show 95% confidence intervals of the pairwise difference between baseline and each subsequent time period.

Center saccade latency also appeared to be unaffected, as there was no statistical evidence for an interaction of Stimulation with Time Period and/or Saccade direction (Table 1). Yet, there was very strong evidence for a main effect of Stimulation in the Bayesian ANOVA. Curiously, this effect was non-significant in the classical ANOVA. This divergence compelled us to delve into the single-subject data, which revealed that one participant showed an effect that was much larger than the other participants (a difference between anodal and cathodal of around 30 ms). When we reran the Bayesian ANOVA without this participant, the inclusion BF_10_ plummeted from 67.1 to 2.4. This participant may have induced a violation of certain assumptions for the Bayesian model, which caused it to behave differently than the classical ANOVA. Still, we decided to run follow-up one-sample tests with this participant included, which showed that latency did not significantly change from baseline for either anodal (*p* = .33, BF_01_ = 3.11) or cathodal (*p* = .41, BF_01_ = 3.52) tDCS. Thus, we conclude that our hypothesis that anodal tDCS would decrease contralateral saccade latency is not supported, and that tDCS had no other effects on saccade latency.

**Table 1:** Classical and Bayesian repeated measures ANOVA results for saccade latency. Factors: *Stimulation* (anodal vs. cathodal), *Time Period* (during tDCS, post-tDCS [0-15 min], post-tDCS [0-30 min]), *Saccade Direction* (left vs. right).

| Effect | df | F | $\eta_G^2$ | p | inclusion $BF_{10}$ | inclusion $BF_{01}$ |
| --- | --- | --- | --- | --- | --- | --- |
| **lateral saccades** |  |  |  |  |  |  |
| Stimulation | 1, 25 | 0.80 | .009 | .38 | 0.76 | 1.32 |
| Stimulation x Direction | 1, 25 | 0.52 | .001 | .48 | 0.22 | 4.56 |
| Stimulation x Time Period | 2, 50 | 0.40 | .0008 | .67 | 0.074 | 13.5 |
| Stimulation x Direction x Time Period | 2, 50 | 2.59 | .003 | .085 | 0.18 | 5.64 |
| **center saccades** |  |  |  |  |  |  |
| Stimulation | 1, 25 | 3.09 | .023 | .091 | 67.2 | 0.015 |

| Effect | df | F | $\eta_G^2$ | $p$ | inclusion BF<sub>10</sub> | inclusion BF<sub>01</sub> |
| --- | --- | --- | --- | --- | --- | --- |
| Stimulation x Direction | 1, 25 | 1.88 | .006 | .18 | 0.74 | 1.34 |
| Stimulation x Time Period | 2, 50 | 0.11 | .0002 | .90 | 0.066 | 15.1 |
| Stimulation x Direction x Time Period | 2, 50 | 1.96 | .001 | .15 | 0.19 | 5.40 |

### Saccade latency distribution

Because the hypothesized effect on median saccade latency was absent, we conducted an additional exploratory analysis (see Quantile analysis in the Material and Methods section) by comparing the entire saccade latency distributions between the anodal and cathodal sessions (Figure 5). Across the board, saccade latencies in the cathodal session were slightly faster than the anodal session, which is opposite to the hypothesized effect of tDCS on FEF excitability. For lateral saccades, the slowest saccades seem to show the biggest difference in latency between the sessions; for center saccades, differences were most pronounced in the fastest saccades. However, these differences were already present in the baseline block, and appear to be driven by a small number of participants. Overall, effects were never significant in the same direction in more than 12 (out of 26) participants.

### Saccade endpoint deviation

No significant effects of tDCS on saccade endpoint deviation were expected, as none were found in Kanai et al. (2012). Yet, at first glance the data seem to show that accuracy improved (i.e. endpoint deviation decreased) with cathodal tDCS (Figure 6). There was a significant and rather large main effect of Stimulation for center saccades, supported by moderate (lateral saccades) to extreme (center saccades) evidence (Table 2). Follow-up one-sample tests for center saccades showed that endpoint deviation only changed significantly from baseline in the cathodal session (*p* = .004, BF_10_ = 10.5), not the anodal session (*p* = .34, BF_01_ = 3.15).

However, this interpretation is muddled by a difference between anodal and cathodal in the baseline, so before tDCS onset (Figure 6). For center saccades, the difference was in fact larger in the baseline than at any other time period (left: mean difference = −0.11°, 95% CI = −0.20° - −0.01°, *p* = .025; right: mean difference = −0.06°, 95% CI = −0.11° - 0.00°, *p* = .066). For example, during tDCS, this difference between anodal and cathodal had completely disappeared (left: *M*_anodal_ = 0.91° = *M*_cathodal_ = 0.91°; right: *M*_anodal_ = *M*_cathodal_ = 0.72°), as endpoint deviation in the anodal session increased from baseline, while it decreased in the cathodal session, thereby cancelling out the baseline difference. Thus, like in Kanai et al. (2012), our results do not appear to support an effect of tDCS on saccade endpoint deviation.

**Table 2:** Classical and Bayesian repeated measures ANOVA results for saccade endpoint deviation. Factors: *Stimulation* (anodal vs. cathodal), *Time Period* (during tDCS, post-tDCS [0-15 min], post-tDCS [0-30 min]), *Saccade Direction* (left vs. right).

| Effect | df | F | $\eta_G^2$ | $p$ | inclusion BF<sub>10</sub> | inclusion BF<sub>01</sub> |
| --- | --- | --- | --- | --- | --- | --- |
| **lateral saccades** |  |  |  |  |  |  |
| Stimulation | 1, 25 | 2.03 | .018 | .17 | 6.64 | 0.15 |
| Stimulation x Direction | 1, 25 | 0.13 | .001 | .72 | 0.19 | 5.21 |
| Stimulation x Time Period | 2, 50 | 0.59 | .002 | .56 | 0.084 | 12.0 |
| Stimulation x Direction x Time Period | 2, 50 | 0.28 | .0003 | .76 | 0.12 | 8.19 |
| **center saccades** |  |  |  |  |  |  |
| Stimulation | 1, 25 | 10.34 | .070 | .004 | 42,209 | 0.00002 |
| Stimulation x Direction | 1, 25 | 2.80 | .013 | .11 | 1.69 | 0.59 |
| Stimulation x Time Period | 2, 50 | 0.61 | .001 | .55 | 0.079 | 12.7 |
| Stimulation x Direction x Time Period | 2, 50 | 0.59 | .001 | .56 | 0.11 | 8.73 |

**Figure 5:**
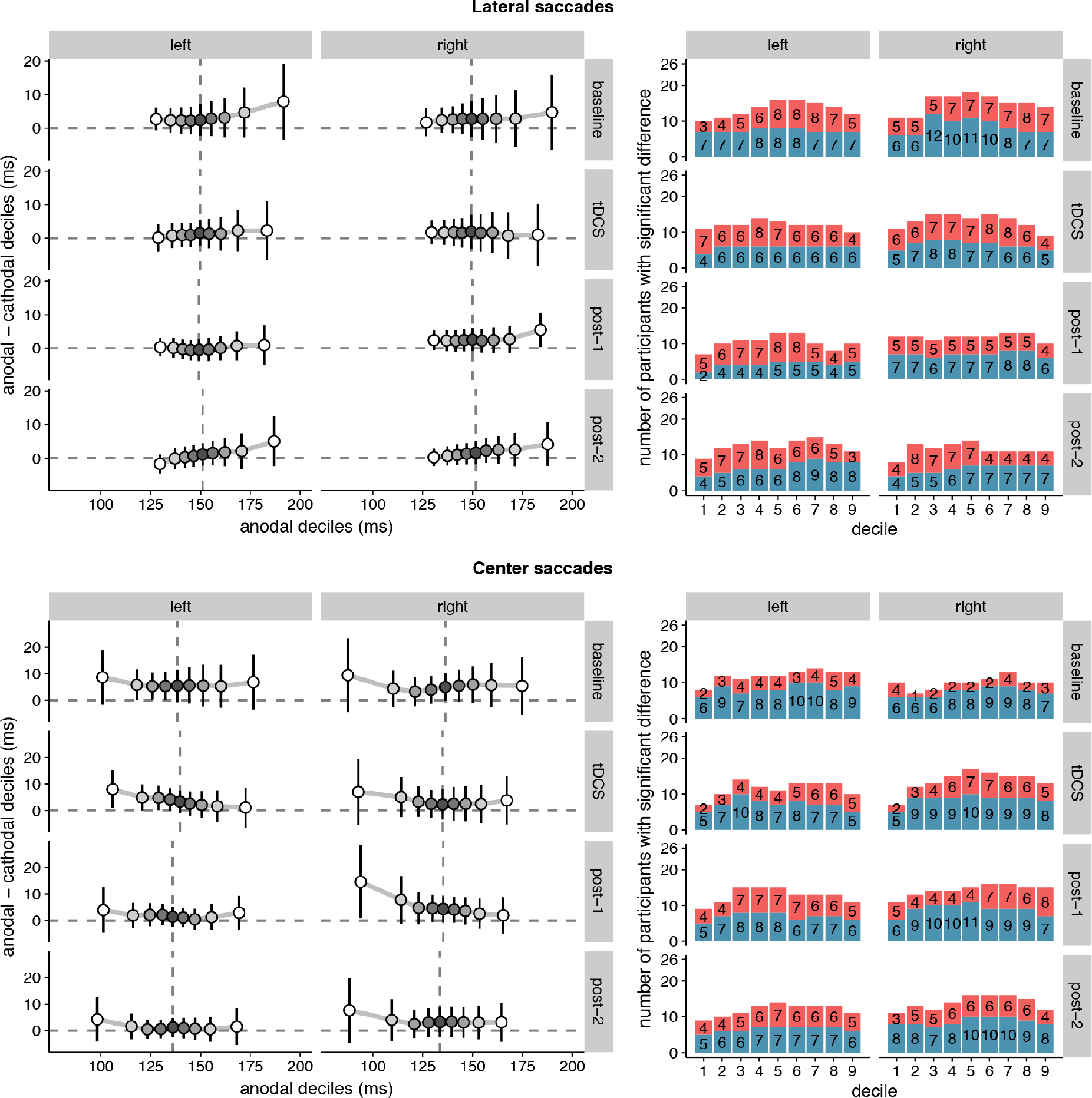
Shift functions of saccade latency distributions under anodal and cathodal tDCS. Data are shown for left- vs. rightward saccades for four 15-minute time periods: baseline, during tDCS, and after tDCS (post-1 and post-2). **Left column**: The x-axis shows saccade latencies for the 9 deciles in the anodal session. The median is plotted as a vertical dashed line. The y-axis shows the difference scores (anodal - cathodal) at each decile. These decile differences express by how much latencies for the cathodal deciles should be *shifted* to match the anodal deciles. Positive differences mean that cathodal saccades had lower latencies than anodal saccades. Error bars show 95% confidence intervals of the decile differences. **Right column**: Counts of participants showing significant effects for the difference between anodal and cathodal sessions at each decile. Red bars count the number of participants with faster anodal saccade latencies; blue bars show counts for faster cathodal latencies. 26 participants is the maximum; the exact number for each contrast is superimposed on the bars.

**Figure 6:**
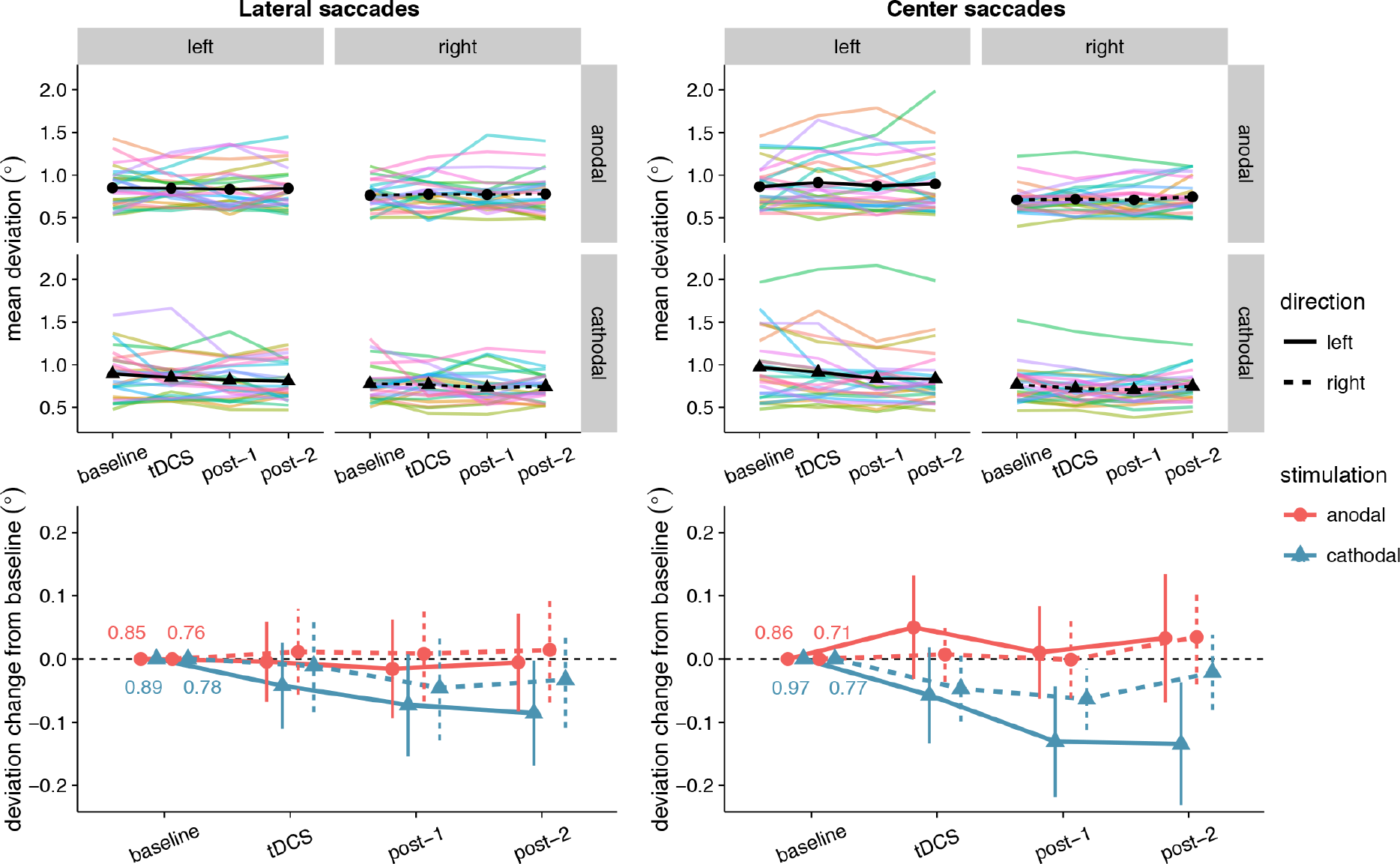
Effects of frontal eye field tDCS on saccade endpoint deviation. Data are shown for left vs. rightward saccades, in the anodal vs. cathodal session, averaged over four 15-minute time periods: baseline, during tDCS, and after tDCS (post-1 and post-2). **Top row**: Colored lines show data from individual participants; black lines show the group median. **Bottom row**: Change in saccade endpoint deviation after baseline subtraction. Numbers inside the plot axes are the baseline saccade endpoint deviations. Error bars show 95% confidence intervals of the pairwise difference between baseline and each subsequent time period.

**Figure 7:**
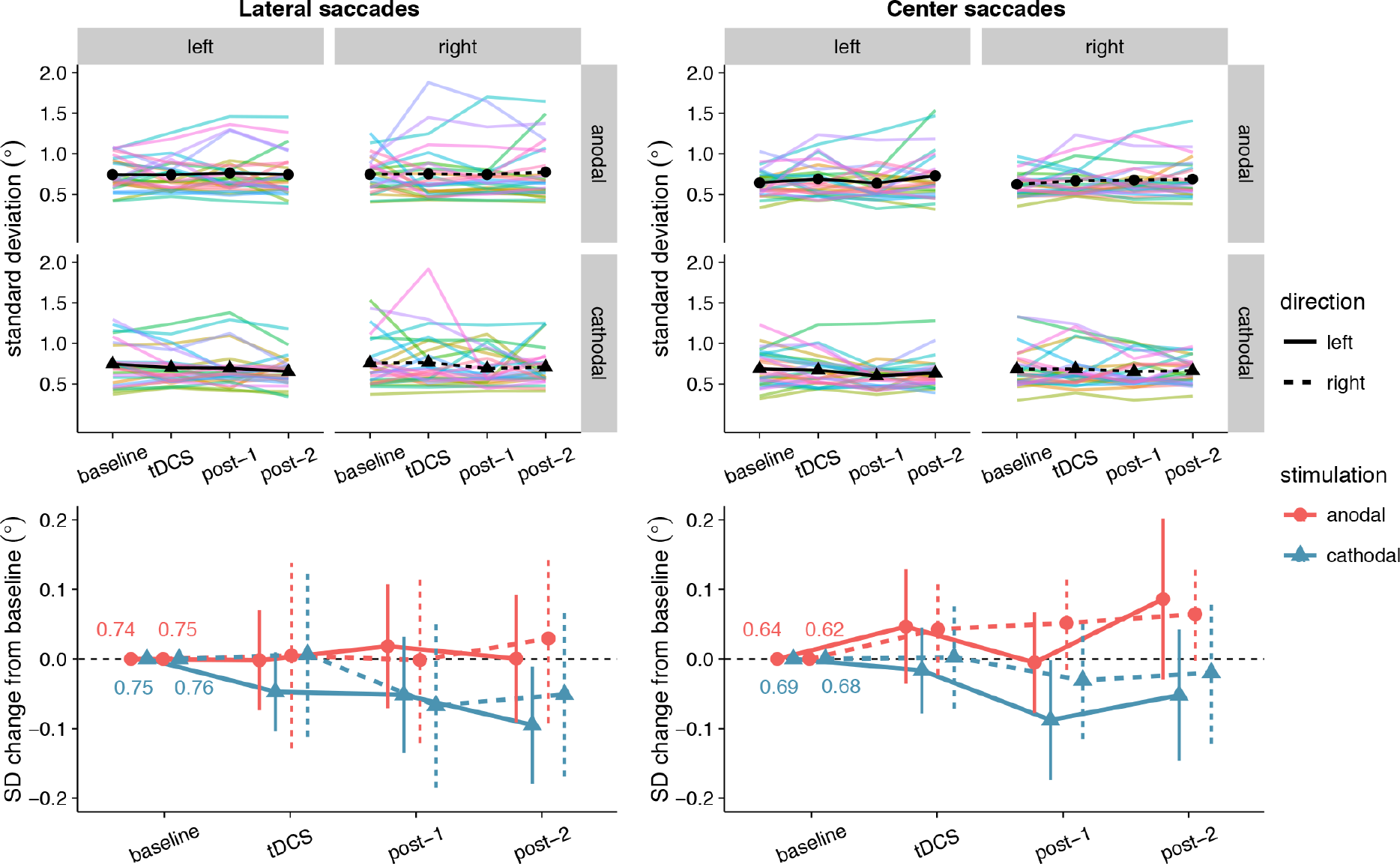
Effects of frontal eye field tDCS on saccade endpoint variability. Data are shown for left vs. rightward saccades, in the anodal vs. cathodal session, averaged over four 15-minute time periods: baseline, during tDCS, and after tDCS (post-1 and post-2). **Top row**: Colored lines show data from individual participants; black lines show the group median. **Bottom row**: Change in saccade endpoint variability after baseline subtraction. Numbers inside the plot axes are the baseline saccade endpoint variability values. Error bars show 95% confidence intervals of the pairwise difference between baseline and each subsequent time period.

### Saccade endpoint variability

Like for saccade endpoint deviation, we had no specific hypotheses on endpoint variability, as Kanai et al. (2012) obtained no effects. However, like the decrease in endpoint deviation, cathodal tDCS also appeared to decrease saccade endpoint variability (Figure 7). For center saccades, there was extreme evidence for inclusion of the main effect of Stimulation in the Bayesian ANOVA, yet the effect only approached significance in the classical ANOVA (Table 3). However, while the variability changes in the anodal and cathodal sessions may have differed from each other, follow-up one sample tests showed that neither anodal (*p* = .11, BF_01_ = 1.40) nor cathodal (*p* = .23, BF_01_ = 2.44) changed significantly from baseline. Thus, saccade endpoint variability also does not seem to be affected by tDCS, conform the findings of Kanai et al. (2012).

**Table 3:**
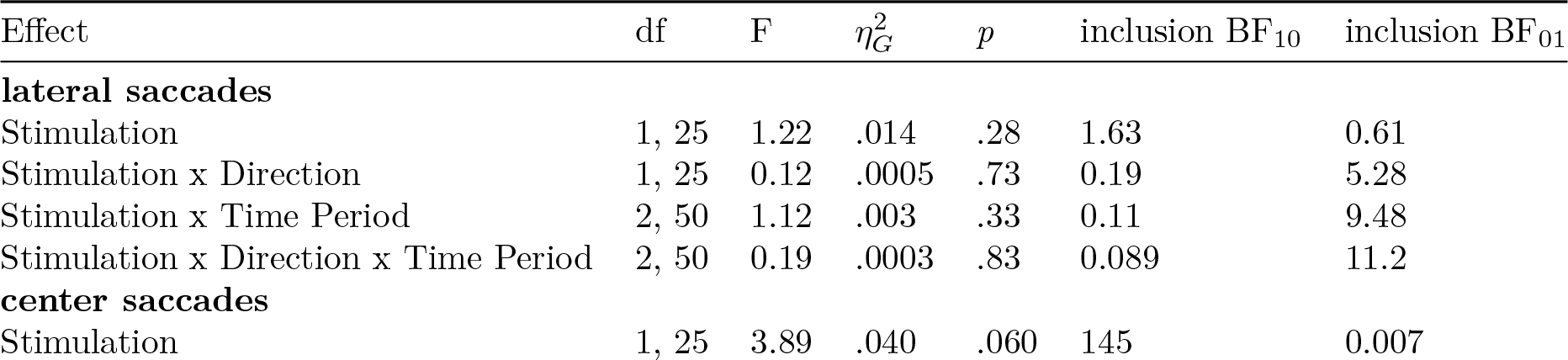
Classical and Bayesian repeated measures ANOVA results for saccade endpoint variability. Factors: *Stimulation* (anodal vs. cathodal), *Time Period* (during tDCS, post-tDCS [0-15 min], post-tDCS [0-30 min]), *Saccade Direction* (left vs. right).

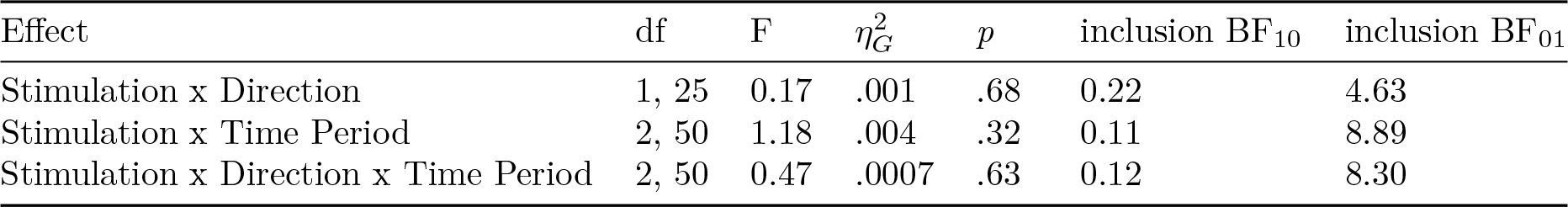

## Discussion

Given the central role the frontal eye field plays in spatial attention, we wanted to examine whether FEF activity could be reliably influenced through tDCS. As the FEF is involved in initiation of eye movements, we measured latency and accuracy of prosaccades to evaluate the effects of tDCS. Our study was based on earlier work (Kanai et al., 2012), which reported that anodal tDCS could speed saccades to targets contralateral to the stimulated FEF. To summarize our results, we were unable to replicate the main effect of Kanai et al. (2012): anodal tDCS did not decrease the latency of contralateral prosaccades. We also found no effects on saccade accuracy, though neither did Kanai et al. (2012).

For saccades back to the center location, Bayesian analyses provided evidence for a differing effect of anodal and cathodal tDCS (regardless of whether saccades were ipsi-/contralateral, or whether they were made during/after stimulation) on all measures we examined: saccade latency, saccade endpoint deviation and saccade endpoint variability. However, in the case of latency and endpoint variability, the corresponding classical analysis was non-significant. Also, follow-up tests (both Bayesian and classical) showed that scores in neither the anodal nor cathodal condition changed significantly from baseline. For endpoint deviation, there was a significant difference between the anodal and cathodal sessions in the baseline, which might have driven the effect. We are therefore hesitant to interpret any of these effects as genuine changes caused by frontal eye field tDCS. Likewise, our shift function analysis painted a complex pattern of differences in the saccade latency distributions for the anodal and cathodal sessions. But these varied highly between individuals and did not seem to exceed the differences that were already present in the baseline block. Collectively, these results do not support an effect of FEF-tDCS on the speed or accuracy of eye movements, and add to a growing body of work that found no results of FEF-tDCS (Ball et al., 2013; Chen and Machado, 2017; Ellison et al., 2014), and tDCS in general (Medina and Cason, 2017; Vöröslakos et al., 2018).

Out of all these results, our null finding for saccade latency is the most surprising. Why did Kanai et al. (2012) find that anodal tDCS speeded (contralateral) saccades, but we did not? Our study should not be considered a direct replication of Kanai et al. (2012), and there are a number of methodological differences between the two. We have tried to enumerate and explain all of them in Table 4. Some are clear and simple improvements, such as the increased statistical power, trial count, and eye tracker sampling rate. Others are more ambiguous: of course, each change was made with the aim to increase the size of the tDCS effect, but each change could also be the cause of why we no longer obtain an effect at all. If that is the case, the changes to the stimulation duration and electrode montage would likely have had the most consequences.

We increased the stimulation duration from 10 to 15 minutes, in order to have more trials during tDCS and possibly a larger neural effect. But longer stimulation durations do not necessarily scale linearly with the effect of tDCS. For example, changing the stimulation duration from 20 to 26 minutes changed the effect of anodal tDCS on motor-evoked potentials from excitatory to inhibitory (Monte-Silva et al., 2013). In addition, we changed the location of the second electrode from the shoulder to the forehead, to more closely resemble the canonical montage used in motor cortex tDCS, and because decreasing the inter-electrode distance can enhance the effect of tDCS (Moliadze et al., 2010). However, next to applying tDCS over the right frontal eye field, it is possible that we now also delivered opposite polarity stimulation to left anterior frontal brain structures. In addition, the exact montage determines to a large extent which brain structures will be in the path of the current—not just those directly under the electrodes, but also those in between (Opitz et al., 2015), as well as distant structures that are anatomically connected (Wokke et al., 2015).

**Table 4:** Methodological differences between the present study and Kanai et al. (2012).

| Difference | Here | Kanai et al. (2012) | Reason |
| --- | --- | --- | --- |
| sample size and design | n=26, within-subject design | n=32, between-subject design | more observations per cell, less influence of between-subject variability |
| FEF localization | MRI-guided per individual | group MRI coordinate | more power (Sack et al., 2009) |
| tDCS: duration | 15 minutes | 10 minutes | more trials during stimulation; possibly increase tDCS effect |
| tDCS: location | right FEF | right or left FEF | right FEF is dominant (Duecker and Sack, 2015) |
| tDCS: montage | FEF, contralateral forehead | FEF, ipsilateral shoulder | decreased interelectrode distance increases effect (Moliadze et al., 2010; Opitz et al., 2015). Resembles canonical motor cortex montage |
| tDCS: conductive medium | Ten20 conductive paste | saline soaked sponges | uniform electrode-skin contact, no risk of excess/leaking saline |
| Number of saccades per condition | 180 (per 15 minutes) | 40 (per 10 minutes) | more robust estimates within each participant |
| Task: stimulus overlap | no overlap of fixation and target | fixation point always on | possibility to analyze saccades back to fixation (center) |
| Task: placeholders | none | target location marked with placeholders | spatial uncertainty might create more room for improvements in accuracy with tDCS |
| Task: ISI | exponential distribution: mean 500 ms, bounds 300-3000 ms | normal distribution, bounds: 300-700 ms | temporally more unpredictable target onsets |
| Eye tracker: sampling rate | 1000 Hz | 250 Hz | more adequate resolution for small effects |
| Eye tracker: saccade threshold | $> 30^\circ/\text{s}$ velocity & $> 8000^\circ/\text{s}^2$ acceleration | $> 26.8^\circ/\text{s}$ velocity | (Eyelink standards) |

We stress that Kanai et al. (2012) also did an experiment with a different electrode montage, in which they delivered bilateral tDCS by placing the anode over the left or right FEF and the cathode over the other FEF (counterbalanced across participants). This montage produced a similar effect, but actually was more effective: tDCS now also speeded saccades contralateral to the anode, but the effect was bigger (7.8 vs. 6.4 ms), and follow-up tests revealed that it was significant at more time points (from 0-30 min after tDCS vs. only 10-20 minutes after tDCS). Nevertheless, we chose to go with a unilateral montage, to be sensitive to possible lateralization of effects. With a bilateral montage, it is impossible to tell whether the effect stems from anodal tDCS to one FEF, cathodal tDCS to the other FEF, or from both at the same time.

Our study was not the first that found no effect of tDCS on saccade latency. Chen and Machado (2017) also set out to replicate this effect, and were also unsuccessful. Like ours, their study was not a direct replication and differed from the protocol used by Kanai et al. (2012) in multiple ways. Specifically, they did not perform MRI-based neuronavigation, and postulated that this might have been the prime reason for why they did not find any effects of tDCS. Although they did place the second electrode on the shoulder, like Kanai et al. (2012), Chen and Machado (2017) also suggest that future studies place it on the left forehead (following the conventional stimulation setup for the motor cortex). Strikingly, our study followed both suggestions (even though our data were collected before their study was published), so it appears these two factors were not responsible for the discrepant results after all.

In addition to the methods, there was also a difference between these studies in average saccade latency. In our study, participants were on average faster (~150 ms for lateral saccades) than in Kanai et al. (2012) (~180 ms). The average center saccade latency was faster still (~135 ms), presumably because the target location was known beforehand in this case. This could be because of changes to the task we made (see Table 4), specifically to have no overlap between target and fixation, and to have no placeholders at the target locations. Both of these are known to reduce saccade latency (Sumner, 2011). Curiously, Chen and Machado (2017) made similar task modifications, but yet obtained not faster but slower saccade latencies (~200 ms).

Although our saccade latencies were faster in the baseline block already, which was thus clearly unrelated to tDCS, this could have diminished the effectiveness of tDCS. The relatively fast latencies could be due to increased inhibition of fixation and an increased proportion of very fast saccades, which rely more on other structures like the superior colliculus (Munoz and Fecteau, 2002; Munoz and Wurtz, 1992) than the fontal eye field. The frontal eye field itself is also not functionally homogeneous: it contains many types of cells (Lowe and Schall, 2017), not just neurons that initiate eye movements, but also those that promote fixation. Even if the frontal eye field was effectively stimulated, there may have been no net effect of tDCS, as the opposing actions of the different cell types could cancel each other out.

Relatedly, the fast saccade latencies could indicate that there was little room for improvement left, and that this task was thus too simple to fully recruit the frontal eye fields. The frontal eye fields are more involved in more effortful tasks, and FEF activity most strongly reflects top-down control (Schafer and Moore, 2011). Lesions of the frontal eye field also impact antisaccades and memory-guided saccades more heavily than simple prosaccades (Rivaud et al., 1994). However, other FEF-tDCS studies that have used more complex tasks like visual search (Ball et al., 2013; Ellison et al., 2017) or the antisaccade task (Chen and Machado, 2017) still found no effects of tDCS.

Perhaps the explanation is also less interesting: the effect could have simply been too small to detect. In Kanai et al. (2012) the reductions in latency produced by tDCS were already fairly modest, especially considering that pioneering studies tend to overestimate effect sizes (Ioannidis, 2008). The neural effects of tDCS itself may also be smaller than anticipated, as recent studies found that the strength of the electric field in the brain is at the lower bound for it to be physiologically effective (Huang et al., 2017; Vöröslakos et al., 2018). As tDCS effects are increasingly viewed as state-dependent, non-linear (Bestmann et al., 2015; Fertonani and Miniussi, 2016) and subject to individual variability (Krause and Cohen Kadosh, 2014; Li et al., 2015), it might be necessary for future studies to use much larger sample sizes (Minarik et al., 2016). It is also vital that future studies employ a sham condition, as there is no a priori guarantee that the anodal/cathodal dichotomy holds for other brain areas (Bestmann and Walsh, 2017) like the frontal eye field.

Such large, well-controlled and more informed (Polanía et al., 2018) studies will be necessary to more clearly establish the boundary conditions of tDCS effects, especially when extending the technique to new brain areas. In the present work, we tried to do so by performing a conceptual replication of the first frontal eye field tDCS study (Kanai et al., 2012). As tDCS did not reliably affect saccade latency or accuracy, we conclude that the efficacy of frontal eye field tDCS remains uncertain.

## Author Contributions

LR and HS conceived the study; LR, TK and HS designed the study. LR and FR collected the data. LR analyzed the data. LR, HS, KR and TK interpreted the data. LR wrote the first draft of the manuscript. LR, TK, KR and HS revised the manuscript. All authors read and approved the submitted version.

## Funding

This work was supported by a VIDI and a Research Talent grant from the Netherlands Organization for Scientific Research (NWO) to HS.

## Acknowledgements

We thank Monja Hoven, Floortje Bouwkamp, and especially Floris Roelofs for their assistance in piloting and data collection. The following colleagues graciously shared their MRI data with us for neuronavigation purposes: Daan van Es, Anouk van Loon, Poppy Watson, Suzanne Oosterwijk, Yaïr Pinto and Henk Cremers.

